# Putative bovine topological association domains and CTCF binding motifs can reduce the search space for causative regulatory variants of complex traits

**DOI:** 10.1101/242792

**Authors:** Min Wang, Timothy P Hancock, Amanda J. Chamberlain, Christy J. Vander Jagt, Jennie E Pryce, Benjamin G Cocks, Mike E Goddard, Benjamin J Hayes

**Affiliations:** Agriculture Victoria, AgriBio, Centre for AgriBioscience, Melbourne, Victoria, Australia; School of Applied Systems Biology, La Trobe University, Melbourne, Victoria, Australia; Faculty of Veterinary and Agricultural Sciences, University of Melbourne, Parkville, Melbourne, Victoria, Australia

**Keywords:** Topological association domains, CTCF binding motifs, allelic-specific expression, allele-specific expression quantitative trial loci, expression quantitative trial loci, functional annotation, cattle

## Abstract

**Background:** Topological association domains (TADs) are chromosomal domains characterised by frequent internal DNA-DNA interactions. The transcription factor CTCF binds to conserved DNA sequence patterns called CTCF binding motifs to either prohibit or facilitate chromosomal interactions. TADs and CTCF binding motifs control gene expression, but they are not yet well defined in the bovine genome. In this paper, we sought to improve the annotation of bovine TADs and CTCF binding motifs, and assess whether the new annotation can reduce the search space for *cis*-regulatory variants.

**Results:** We used genomic synteny to map TADs and CTCF binding motifs from humans, mice, dogs and macaques to the bovine genome. We found that our mapped TADs exhibited the same hallmark properties of those sourced from experimental data, such as housekeeping gene, tRNA genes, CTCF binding motifs, SINEs, H3K4me3 and H3K27ac. Then we showed that runs of genes with the same pattern of allele-specific expression (ASE) (either favouring paternal or maternal allele) were often located in the same TAD or between the same conserved CTCF binding motifs. Analyses of variance showed that when averaged across all bovine tissues tested, TADs explained 14% of ASE variation (standard deviation, SD: 0.056), while CTCF explained 27% (SD: 0.078). Furthermore, we showed that the quantitative trait loci (QTLs) associated with gene expression variation (eQTLs) or ASE variation (aseQTLs), which were identified from mRNA transcripts from 141 lactating cows’ white blood and milk cells, were highly enriched at putative bovine CTCF binding motifs. The most significant aseQTL and eQTL for each genic target were located within the same TAD as the gene more often than expected (Chi-Squared test P-value ≤ 0.001).

**Conclusions:** Our results suggest that genomic synteny can be used to functionally annotate conserved transcriptional components, and provides a tool to reduce the search space for causative regulatory variants in the bovine genome.

## Background

Identifying causal mutations is essential for improving the accuracy and reliability of genomic selection [1]. This identification task is challenging because the large scale linkage disequilibrium and small effects of most mutations in the bovine genome can drive a false discovery in a genome-wide association study (GWAS) [2]. To distinguish the causal mutations from noise, gene expression can be utilised to identify genomic loci that shape the trait of interest. An expression quantitative trait locus (eQTL) is a heterozygous locus that is associated with total changes in a gene’s expression in a group of individuals. An allele-specific expression quantitative trait locus (aseQTL) is a heterozygous locus that explains allele-specific expression of a particular gene transcript in a group of individuals. Here, the heterozygous locus is called eSNP or eVariant, and the gene that displays expression variation is called eGene [3–7]. Both aseQTL and eQTL mappings require high computational capacities to test association between any eSNP and any eGene genome-wide. To reduce the computing time and space, we propose to take into account the transcriptional regulatory structures that confine the scope of chromosomal interactions, and only test *cis*- association between eSNPs and eGenes under the same transcriptional control.

Topological association domain (TAD) is a type of regulatory structure that has never been described in the bovine genome. Topological association domains (TADs) are empirically defined from Hi-C or other chromatin conformation data [8–13]. TADs partition the genome by contact frequencies, where chromosomal regions within the same domain self-interact much more frequently than between domains. TADs are highly conserved between cell types and species [8, 14–16], and the disruptions of TADs were found to cause disease-related gene expression by exposing genes to inappropriate regulatory elements [13, 17, 18]. A TAD boundary is defined as genomic interval, no larger than 400 Kb, between adjacent TADs where self-interactions decrease [14]. The TAD boundaries were found to be enriched for housekeeping genes, transfer RNA (tRNA) genes, tri-methylation of lysine 4 on histone H3 (H3K4me3) signals, short interspersed nuclear elements (SINEs), DNase hypersensitive sites, and CTCF binding sequences [9, 14, 19, 20].

The CCCTC-binding factor (CTCF) is an 11 zinc-finger protein that mediates transcriptional regulation. CTCF can bind to evolutionarily conserved DNA sequences to prevent inappropriate enhancer-promoter interactions, and these conserved CTCF binding sequences were found to be enriched at TAD boundaries [14, 21]. CTCF can also bind to evolutionarily non-conserved DNA sequences to facilitate unique enhancer-promoter interactions at various steps of the transcriptional process, and these diverse CTCF binding sequences are distributed within TADs [14, 22, 23]. The CTCF protein has a highly conserved amino acid sequence that is 93% identical from avian to human, and its functional diversities are achieved through different combinations of zinc finger domains binding to DNA sequences [24]. These DNA sequences have distinctive patterns, called CTCF binding motifs, and some of them are highly conserved across 180 million years of evolution [25]. Similar to TADs, mutations at CTCF binding motifs are commonly deleterious [26].

In this study, we aimed to assist in reducing the search space for causative regulatory variants by identifying TADs and CTCF binding motifs in the bovine genome. We mapped our bovine TADs and CTCF binding motifs based on homology with humans, mice, dogs and macaques. Our bovine TADs and CTCF binding motifs were highly enriched for well-known biological hallmarks. We subsequently showed that our new bovine TADs and CTCF binding motifs could predict runs of genes that were reported by Chamberlain *et al*. (2015) [27] as favouring the same parental allele. Finally, we showed that aseQTLs and eQTLs, which were identified from mRNA transcripts from 141 lactating cows’ white blood and milk cells [28], were enriched at putative CTCF binding motifs, and were located in the same TAD as their eGene target more often than expected. This finding, where mRNA expression profiles across multiple lactating cows’ tissues and cells were confined within TADs and CTCF binding motifs, indicated that our putative TADs and CTCF binding motifs could be used to reduce the search space for causative regulatory variants in the bovine genome.

## Results

### Mapping mammalian topological association domains to the bovine genome

We have directly mapped large genomic segments, the topological association domains (TADs), to the bovine genome, and tracked the changes in each step of the mapping. We found that when genomic conversion was to an updated version of reference assembly of the same species (hg18 to hg19), the query TADs recovered well in the target genome (S1 Table). Over 89% of query TADs mapped uniquely to a single location in the target reference assembly without splitting within (intra-chromosomal) or across (inter-chromosomal) chromosomes. The query and target TADs not only had similar widths, but also had little variation in genomic positions. When the genomic conversion was to an older reference assembly of the same species (e.g. canfam3 to canfam2), the query TADs recovered less well. Less than 10% of query TADs mapped uniquely to a single location in the target genome, although over 76% were intra-chromosomal splits. When genomic conversion was across species (hg19 to bostau6, mm9 to bostau6, mm10 to bostau7 and canfam2 to bostau6), over 98% of query TADs split and over 87% were inter-chromosomal splits. In the most extreme case, a query TAD was broken down into 157,709 locations across multiple target chromosomes. No input TADs aligned to chromosome Y or mitochondrial chromosome, but there were TAD fragments that aligned to the bovine mitochondrial chromosome across all cohorts of input TAD sets. Overall, the TADs from human and dog mapped better to the bovine genome than those from mouse, likely due to a closer evolutionary distance among human, dog and bovine than between mouse and bovine.

The mapping of large genomic segments inevitably created a large number of genomic fragments. We mended TAD fragments via a recovery, a filtering and a local refinement procedure. Our method resulted in putative bovine TADs that resembled the respective input TADs in the following aspects: firstly, similar number of TADs were found in the bovine genome as the respective input TADs (Table 1); secondly, no putative bovine TADs aligned to chromosome Y or mitochondrial chromosome (S1 Table); thirdly, TAD widths were comparable between bovine and the source TADs (Fig 1); finally, the majority of input TADs were mapped to the bovine genome (79.63-98.88%), and 65.55-89.03% of those mapped uniquely to a single genomic location without any intra-chromosomal or inter-chromosomal splits (Table 1).

**Table 1.**
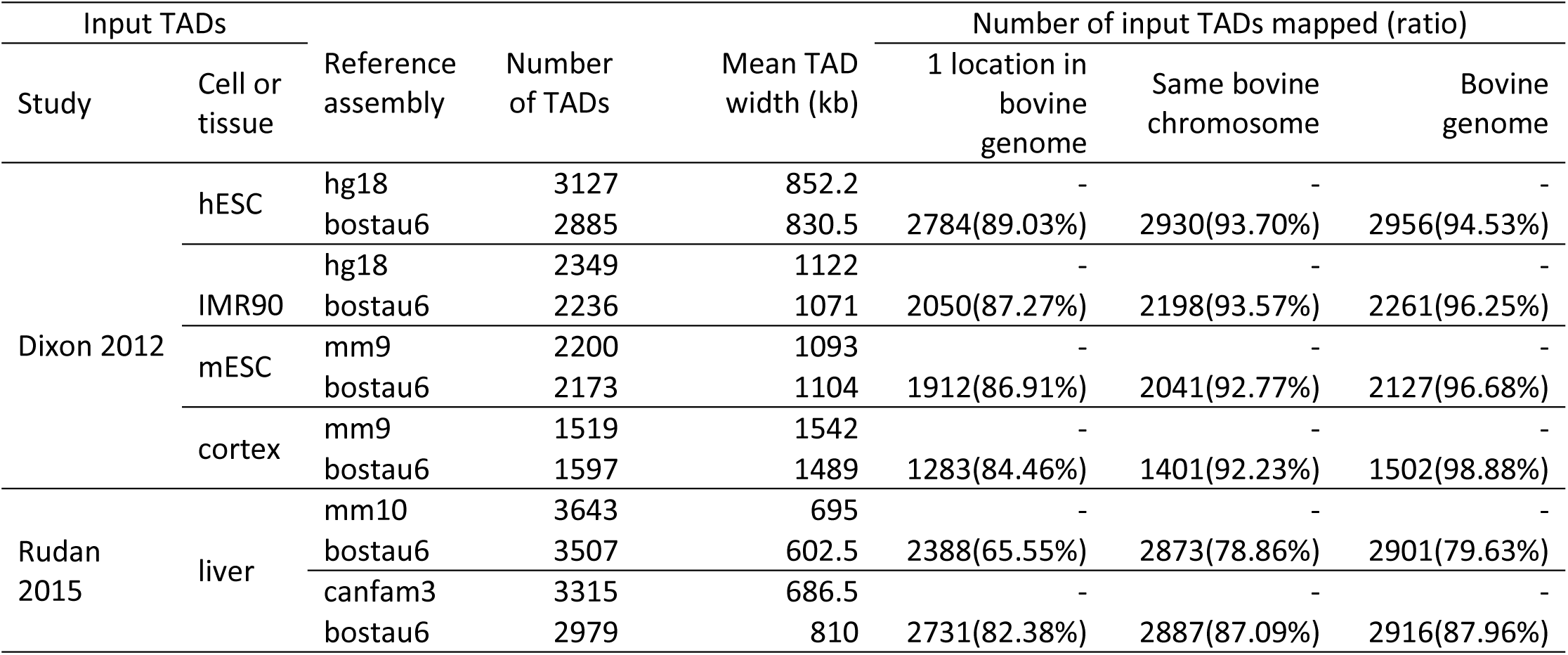
Summary statistics of TAD mapping. The source study and cell/tissue from each input dataset, and also the input and output reference assemblies with the number of TADs and their mean width (in kilobases) are shown. Also presented are the number of input TADs mapped to a single location in the bovine genome, the same bovine chromosome and the bovine genome.

**Fig 1.**
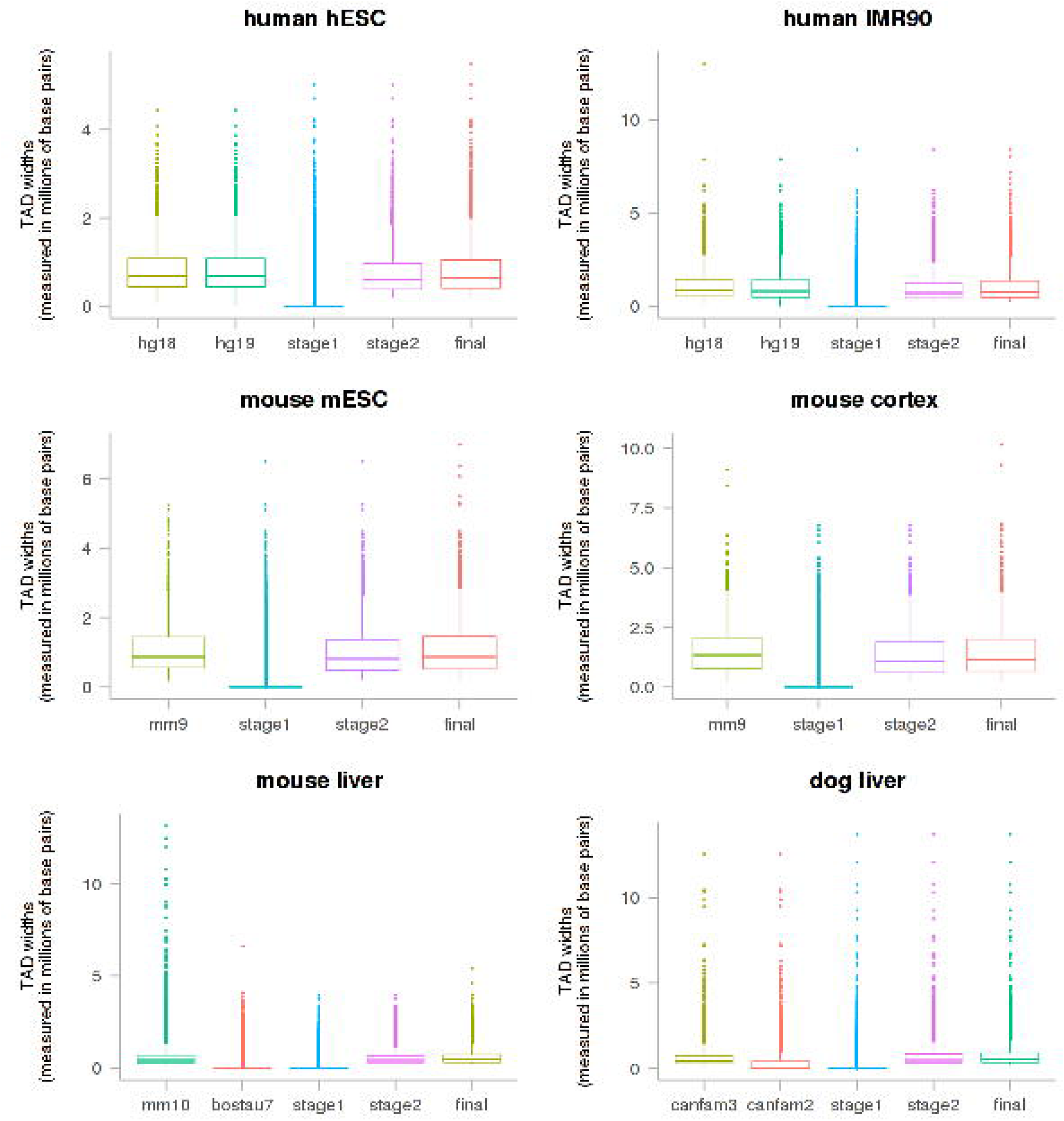
Distribution of TAD widths. For each input dataset, the TAD width (millions of base pairs) of the input data, intermediate reference (if it was used), putative bovine TADs (stage 1 and stage 2) and final putative bovine TADs were shown as boxplots.

### Scanning for putative bovine liver CTCF binding motifs

Chromatin immunoprecipitation with sequencing (ChIP-Seq) is an assay that identifies where a protein and DNA interact in the sample. We downloaded a total of 184,492 CTCF ChIP-Seq data, which ranged from 42 bp to 1,716 bp from the liver tissue of human, mouse, dog and macaque (Table 2). We found a total of 82 clusters of motif profiles within all those CTCF ChIP-Seq data. Of those, 4 clusters that were validated in the JASPAR, UniProt or HOCOMOCO databases (S1 Appendix) were widely reported in literature [16, 24–26, 29–31]. All 82 clusters of motifs were scanned across bovine chromosome 1 to X, and a total of 3,770,311 putative bovine CTCF binding motifs were found (P-value ≤ 10^−5^; S1 Appendix). Note that if there were 2 motifs on the same genomic region but were respectively on the forward and reserve DNA strands, they were counted as 2 motifs. The putative bovine CTCF binding motifs were short (7-29bp) and tended to group into clusters distributing sparsely across the entire bovine genome. Less than 14% putative bovine CTCF binding motifs overlapped with another putative bovine CTCF binding motif. We defined a more restrictive set of CTCF binding motifs as the putative bovine CTCF binding motifs whose motif score was no less than 80, motif P-value was no larger than 10^−8^, and any overlapping regions on the same DNA strand were merged. We found 78,524 more restrictive CTCF binding motifs on the bovine genome. We defined CTCF gaps as the genomic intervals between the more restrictive CTCF binding motifs. There were 45,809 CTCF gaps ranging from 1 bp to 1,503,697 bp.

**Table 2.**
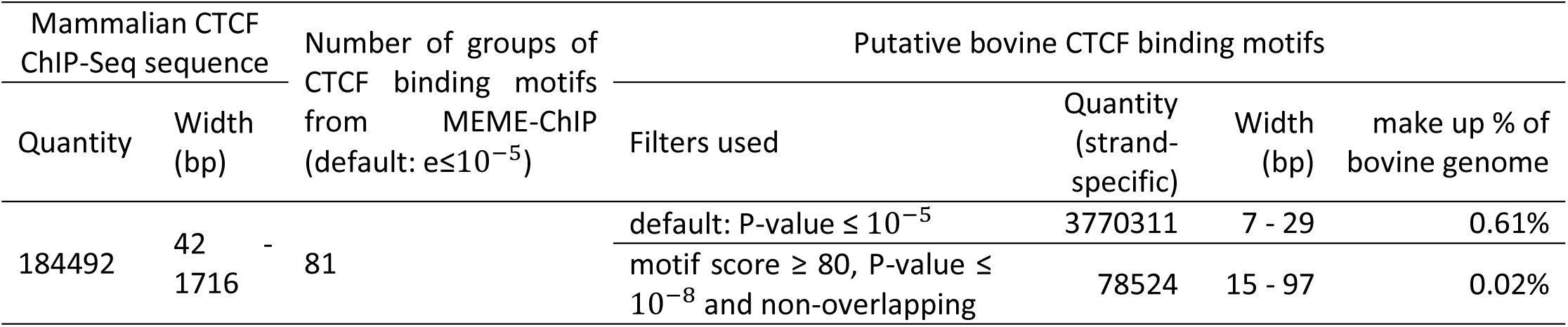
Properties of putative bovine CTCF binding motifs and selected putative bovine CTCF binding motifs

### Enrichment of biological hallmarks at TAD boundaries

We validated our final sets of bovine TADs by assessing the level of enrichment of biological hallmarks at the TAD boundaries (Fig 2; S2 Table A). We found that putative bovine CTCF binding motifs (P-value ≤ 10^−5^), bovine liver H3K27ac and H3K4me3 regions, bovine SINE and tRNA genes were all highly enriched in all sets of bovine putative TADs. House-keeping genes were highly enriched in all sets of bovine putative TADs but less so in putative bovine TAD set mapped from mouse liver. Bovine H3K4me3 regions from tender and tough muscle tissues were highly enriched in all sets of bovine putative TADs but less so in putative bovine TAD sets mapped from mouse cortex and liver. Putative bovine enhancers in homology with VISTA, FANTOM5 and dbSUPER datasets were not enriched in all sets of TAD boundaries. The same permutation test was repeated but excluding ‘knots’, which meant that we assumed there were no TAD boundaries if the end position of the previous TAD was the same as the start position of the following. We found a very similar enrichment profile as those including knots (S2 Table A). The same permutation test was also repeated using the more restrictive set of CTCF binding motifs (motif score ≥ 80 and motif P-value ≤ 10^−8^), but we did not see significant enrichment of the more restrictive set of CTCF binding motifs at TAD boundaries across all TAD sets except those mapped from the mouse liver TAD set (S2 Table A).

**Fig 2.**
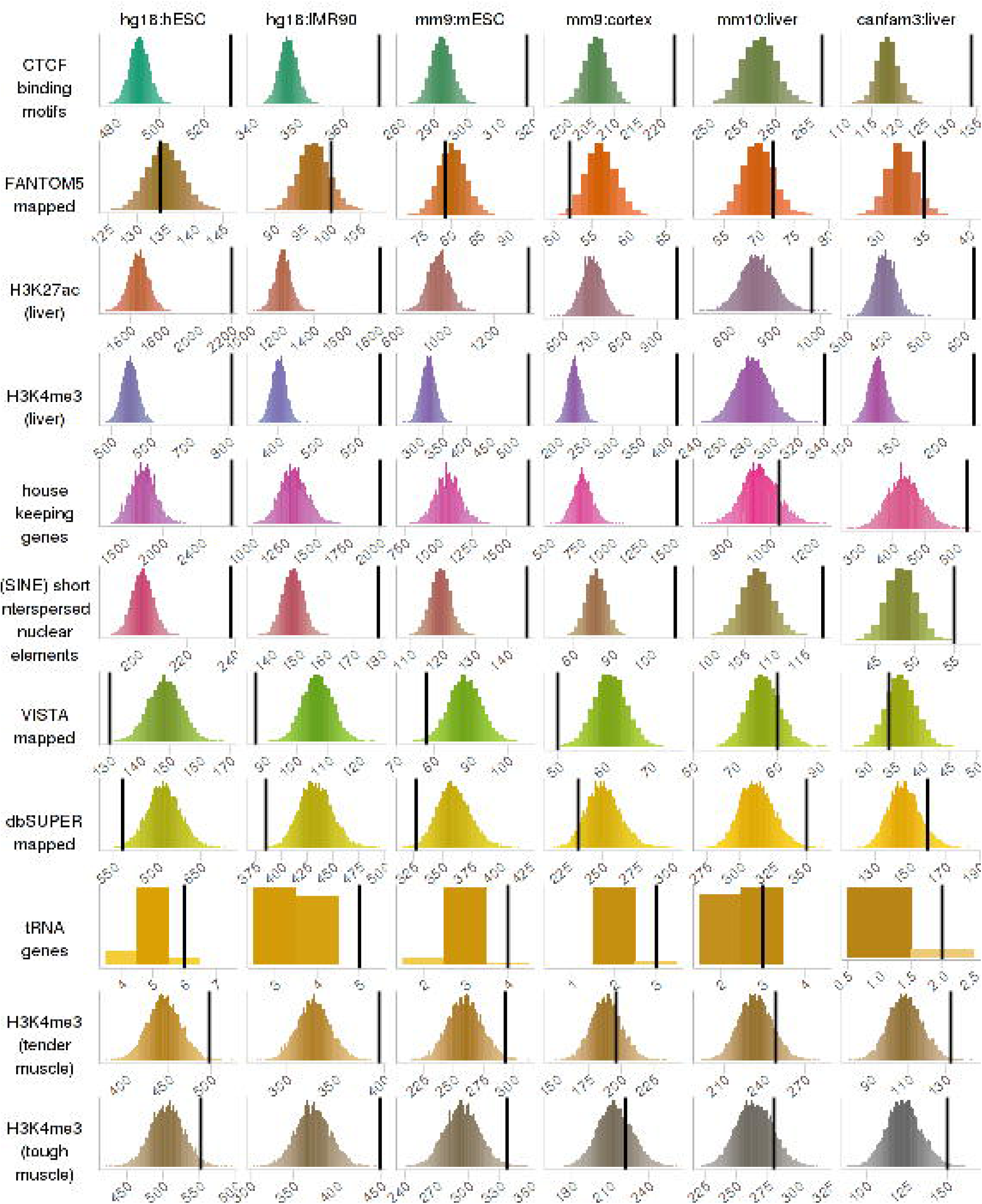
Enrichment of biological hallmarks in putative bovine TAD boundaries. For each hallmark biological signal (columns) and each putative bovine TAD set (rows), these frequency histograms show the number of overlapping base pairs between the signal and the TAD boundaries. The frequencies were rounded to the nearest 10 thousands, and the bin width was 1. The 10,000 random permutations are in colour and the actual number is the black vertical line. If a biological signal is enriched at TAD boundaries, the vertical line will be on the right and clearly separated from the histogram.

A total of 108 types of CTCF binding motifs were found in the bovine genome (S1 Appendix). To examine which CTCF binding motifs were enriched at TAD boundaries, we repeated the same enrichment analysis for each motif pattern (S2 Table B). We found that the four motif patterns that were validated in public databases were all highly enriched at all sets of putative bovine TAD boundaries (S1 Appendix; S2 Table). Some putative bovine CTCF binding motifs, even though were not validated by public transcriptional factor binding motifs databases, were highly enriched at all sets of TAD boundaries (e.g. motif number 15 and 77 in S2 Table B). Some putative bovine CTCF binding motifs were only enriched at putative bovine TAD boundaries mapped from cell lines but were not as enriched at putative bovine TAD boundaries mapped from tissues (e.g. motif number 6 and 11 in S2 Table B). A large number of motifs from the less restrictive set of putative bovine CTCF binding motifs (P-value ≤ 10^−5^) were absent in the more restrictive set of putative bovine CTCF binding motifs (motif score ≥ 80 and motif P-value ≤ 10^−8^), but only a few of those were enriched at TAD boundaries (e.g. motif number 52 and 77 in S2 Table B), and the majority were not enriched.

### Testing allele-specific expression within regulatory units

Runs of genes were found to favour the same parental allele in 1 lactating cow’s 18 tissues [27]. We used analysis of variance (ANOVA) to test whether exons within any regulatory units were significantly biased for expressing from a parental chromosome in a tissue. The regulatory units were respectively defined only by TAD, only by the more restrictive set of CTCF binding motifs, and by both TAD and more restrictive set of CTCF binding motifs in three ANOVA models. Of those 108 cohorts (6 TAD sets × 18 tissues) tested in the TAD only model, we found that on average 14% of allele-specific expression (ASE) variation were explained, with a standard deviation (SD) of 0.056 across 107 significant cohorts (P ≤ 10^−8^). The remaining cohort, putative bovine TAD set from dog liver, did not show significant ASE variation in lung. The putative bovine TADs from the mouse cortex explained the least amount of ASE variation in white skin (3.45%), and those from mouse liver explained the largest amount of ASE variation in white blood cells (28%; Fig 3). The CTCF only model had 6% less ASE SNPs than the TAD only model had, and explained on average 27% (SD: 0.078) ASE variation across all 18 significant cohorts (1 CTCF gaps × 18 ASE tissues; P ≤ 10^−8^). The CTCF only model explained ASE variation across 18 tissues in a trend that was similar to the TAD only model, where the least amount of ASE variation explained was in lung (10%) followed by liver (17%), and the largest amount of ASE variation explained was in white blood cells (42%; Fig 3). The TAD+CTCF model tested around 30% less ASE SNPs than the TAD only model, and explained on average 31% (SD: 0.089) ASE variation across all 108 significant cohorts (6 TAD sets × 18 tissues; P ≤ 10^−8^). Trend was also similar to the TAD only model and CTCF only model, where the least amount of ASE variation explained was also in lung (on average 11%; SD: 0.012), and the largest amount of ASE variation explained was also in white blood cells (on average 48%; SD: 0.023; Fig 3). We found little increments, and sometimes even a slight decrease, in variation explained by the TAD+CTCF model in comparison to the CTCF only model, which indicated that the CTCF model encompassed the TAD model, and was better at explaining ASE variation. Our permutation tests declared strong significance for all cohorts in all models tested, meaning that our observation that the confinement of ASE variation within regulatory units was not random.

**Fig 3.**
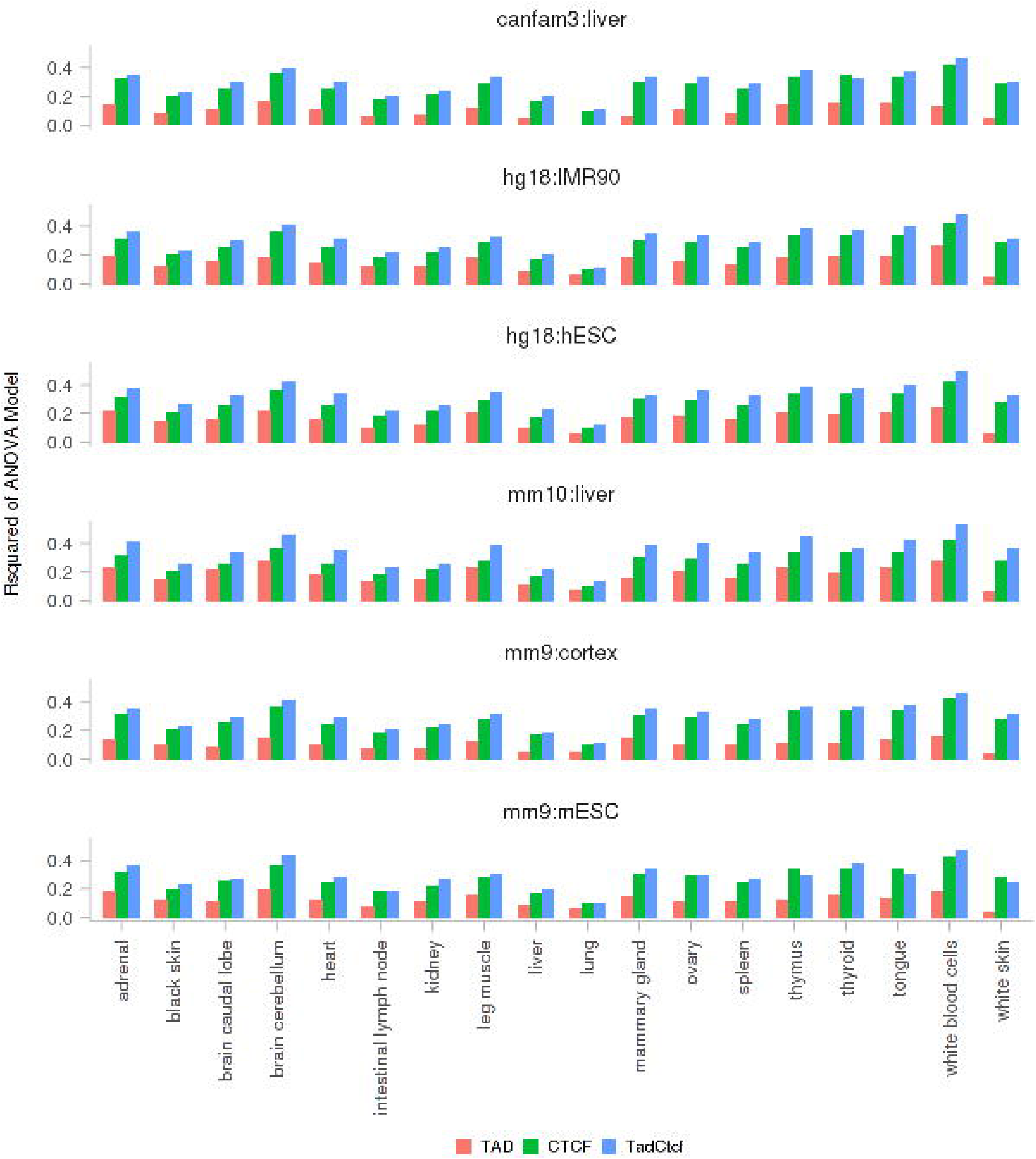
R-squared values from significant ANOVA models (P ≤ 10^−8^). For each tissue (X-axis) and each regulatory unit defined by TAD, CTCF gaps and TAD+CTCF gaps, these bar plots show the R-squared values of significant (P ≤ 10^−8^) ANOVA models (Y-axis). Note in the CTCF model, there is no input TAD set involved. For the purpose of a convenient comparison with the other models, the R-squared values in the CTCF model were plotted across all cohorts from the same tissue.

### Enrichment of significant aseQTLs and eQTLs within putative bovine CTCF binding motifs

We defined significant aseQTLs and eQTLs by the P-value from aseQTL and eQTL mappings respectively [28]. We found across all P-value thresholds tested ranging from 10^−5^ to 10^−8^, the significant aseQTLs and eQTLs in white blood and milk cell were highly enriched at putative bovine CTCF binding motifs (P-value ≤ 10^−5^; Table 3) in comparison to the null distribution sampled from the entire bovine genome. The averaged fold change of enrichment for significant aseQTLs in white blood cells was 1.55 fold and in milk cells was 1.49 fold. The averaged fold change of enrichment for significant eQTLs in white blood cell was 1.43 fold and in milk cells was 1.41 fold (Table 3). The same permutation test was repeated in the more restrictive set of CTCF binding motifs (motif score ≥ 80 and motif P-value ≤10^−8^), and significant aseQTLs and eQTLs in white blood and milk cell were also highly enriched across all P-value thresholds tested (Table 3; Fig 4). The fold change of enrichment was higher in the more restrictive set of CTCF binding motifs. On average, the significant aseQTLs in white blood cells were 4.06 fold and in milk cells were 3.74 fold, and the significant eQTLs in white blood cells were 3.23 fold and in milk cells were 4.18 fold more enriched in comparison to the rest of the genome (Table 3; Fig 4). The same method was also used to test the level of enrichment for significant aseQTLs and eQTLs in biological hallmarks including H3K4me3 regions from both bovine liver and muscle tissues, and H3K27ac regions from bovine liver tissue, and bovine genomic regions in homology with VISTA, FANTOM5 and dbSUPER databases, but none of those biological signals were as enriched for significant aseQTLs and eQTLs as the bovine putative CTCF binding motifs did across all P-value thresholds tested (S3 Table).

**Table 3.** Rank and fold change of enrichment of significant aseQTLs and eQTLs at putative bovine CTCF binding motifs

| QTL type | cell type | P-value significant threshold | motif P-value $\leq 10^{-5}$ | | motif score $\geq 80$ and motif P-value $\leq 10^{-8}$ | |
| --- | --- | --- | --- | --- | --- | --- |
|  |  |  | rank | fold change | rank | fold change |
| aseQTL | white blood | $10^{-5}$ | <0.0001 | 1.54 | <0.0001 | 4.19 |
| | | $10^{-6}$ | <0.0001 | 1.55 | <0.0001 | 4.07 |
| | | $10^{-7}$ | <0.0001 | 1.55 | <0.0001 | 4.02 |
| | | $10^{-8}$ | <0.0001 | 1.56 | <0.0001 | 3.96 |
| | milk | $10^{-5}$ | <0.0001 | 1.49 | <0.0001 | 3.97 |
| | | $10^{-6}$ | <0.0001 | 1.49 | <0.0001 | 3.71 |
| | | $10^{-7}$ | <0.0001 | 1.49 | <0.0001 | 3.67 |
| | | $10^{-8}$ | <0.0001 | 1.48 | <0.0001 | 3.61 |
| Eqtl | white blood | $10^{-5}$ | <0.0001 | 1.38 | <0.0001 | 3.97 |
| | | $10^{-6}$ | <0.0001 | 1.43 | <0.0001 | 3.56 |
| | | $10^{-7}$ | <0.0001 | 1.45 | <0.0001 | 2.75 |
| | | $10^{-8}$ | <0.0001 | 1.47 | <0.0001 | 2.66 |
| | milk | $10^{-5}$ | <0.0001 | 1.34 | <0.0001 | 4.48 |
| | | $10^{-6}$ | <0.0001 | 1.40 | <0.0001 | 4.75 |
| | | $10^{-7}$ | <0.0001 | 1.38 | <0.0001 | 4.05 |
| | | $10^{-8}$ | <0.0001 | 1.53 | 0.9978 | 3.44 |

**Fig 4.**
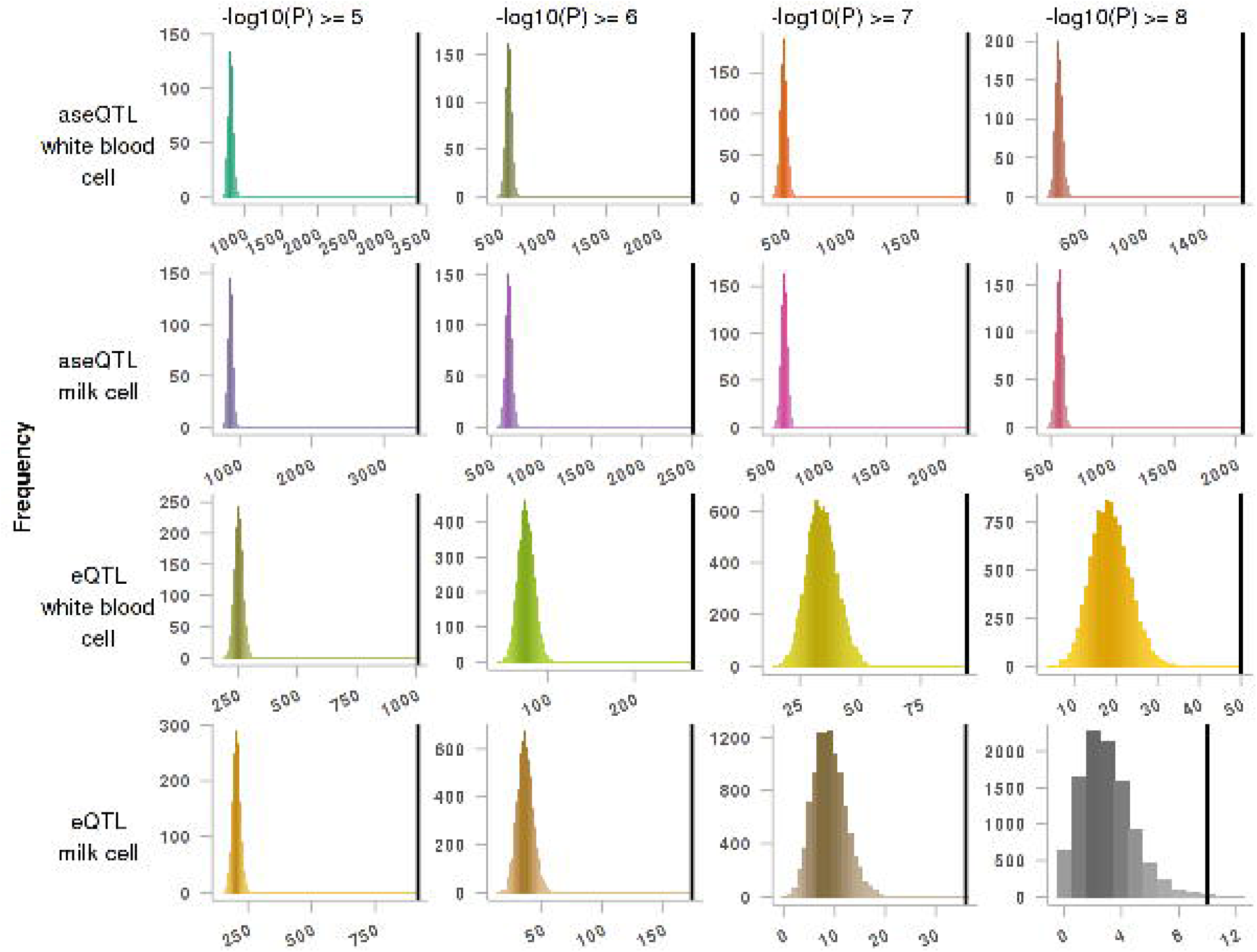
Enrichment of significant aseQTLs and eQTLs in putative bovine CTCF binding motifs. For each significant threshold (columns) and each aseQTL/eQTL in white blood cells or milk cells (rows), these frequency histogram show the number of significant aseQTLs/eQTLs in putative bovine CTCF binding motifs (motif score ≥ 80 and motif P-values ≤ 10^−8^). The 10,000 random permutations are in colour and the actual number is the black vertical line. If the actual significant aseQTLs/eQTLs are enriched at putative bovine CTCF binding motifs, the vertical line will be on the right and clearly separated from the histogram.

### Testing frequency of the most significant aseQTL or eQTL within the same TAD as eGene target

We required the most significant aseQTL and eQTL for each eGene to pass a P-value threshold, which was tested from 10^−5^ to 10^−8^. When multiple aseQTLs or eQTLs were ranked the highest significance towards the same eGene, we selected linearly the furthermost aseQTL and eQTL for each eGene in order to avoid bias favouring the aseQTL and eQTL in a linearly closer distance to the eGene. Across all P-value thresholds tested and in both white blood and milk cell samples, the observed numbers of furthermost, and most-significant, aseQTLs and eQTLs within the same TAD as their eGene were all significantly more than expected, with the Chi-Squared test P-values all close to 0 (Table 4; S4 Table). This indicated that our putative bovine TADs could provide the right space to search for *cis*-regulatory variants and their genic targets.

**Table 4.**
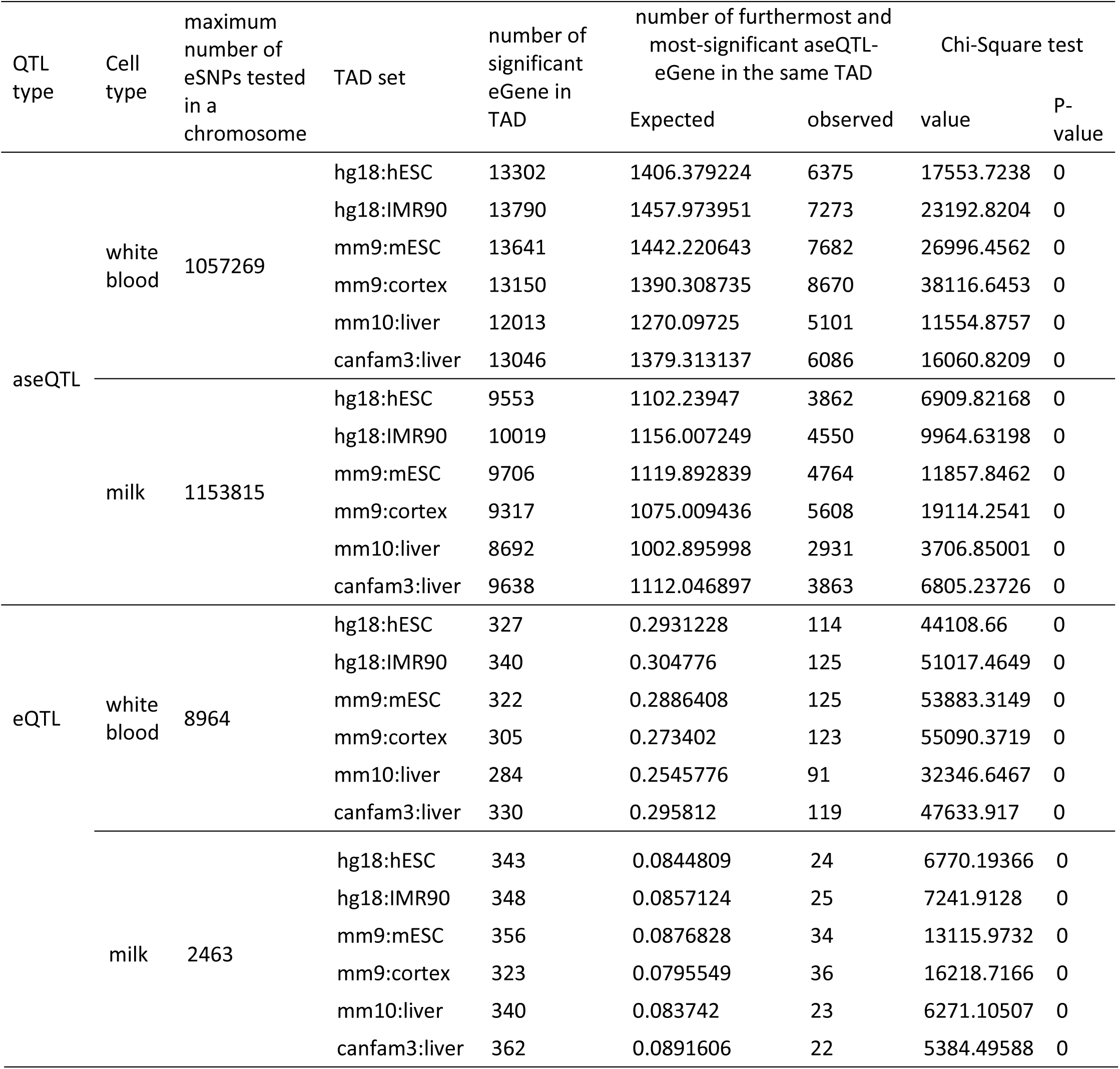
eGenes are often located within the same TAD as their furthermost and most-significant aseQTL/eQTL. The most-significant aseQTL and eQTL for each eGene all have a P-value less than 10^−8^ in aseQTL/eQTL mapping.

## Discussion

We have presented a computational method that identifies topological association domains (TADs) based on sequence homology. Our method relies on the conserved nature of TADs, the evolutionary distance between species and the quality of reference genome assemblies. A small proportion (1%-21%) of mammalian TADs were filtered out in the final bovine TAD sets (S1 Table), because they did not fulfil the size (>200Kb) or consensus (>3 TAD sets agreed) requirement. Input TADs from species evolutionarily closer to bovine were mapped better than those further from bovine (S1 Table). Input TADs from the latest version of reference assemblies were mapped better than those from older version of reference assemblies of the same species (S1 Table). Overall, our mapping results agree with observations that TADs are largely invariant in development and evolution [14, 16, 32, 33]. Most input TADs were intact in the bovine genome (Table 1; S1 Table). Mammalian TADs from different input TAD sets were mapped to similar locations in the bovine genome. Our bovine TADs also displayed key features of mammalian TADs from HiC data (Fig 1; Fig 2; S1 Table), indicating that our TADs were a good proxy for the actual TADs in the bovine genome.

We have also created a library of putative bovine CTCF binding motifs from mammalian CTCF ChIP-Seq data, due to the absence of bovine CTCF binding sequences. Only a small proportion of putative bovine CTCF binding motifs was previously reported, and was validated by transcription factor binding motifs that were made available to public databases (S1 Appendix). The validated CTCF binding motifs tended to be highly conserved and were highly enriched at all sets of TAD boundaries (S2 Table). However, our putative bovine CTCF binding motifs were validated by different databases but not all, and some non-validated CTCF binding motifs were also enriched at all sets of putative bovine TAD boundaries. These indicate that the current CTCF binding motifs in public databases were incomplete, and different public motif databases complement each other. If more known motifs were supplied for validation, more putative bovine CTCF binding motifs could be validated.

To compensate the incomplete validation from motif databases, we compared results from two sets of putative bovine CTCF binding motifs that were differed by P-value stringencies. The less restrictive set of putative bovine CTCF binding motifs (motif P-value ≤ 10^−5^) were more enriched at TAD boundaries than the more restrictive set (motif score ≥ 80 and motif P-value ≤ 10^−8^; S2 Table). One explanation for this observation is that the more restrictive set of putative bovine CTCF binding motifs were too sparsely distributed across the bovine genome and therefore failed an enrichment analysis. We observed where the more restrictive set of CTCF binding motifs were found, similar motif patterns were also found in the less conserved set (S2 Table). This implies the same genomic locations tagged.

TADs are defined from Hi-C data which snapshots chromosomal regions in close proximities without specifying any target loci [8, 10]. Contact maps similar to TAD can also be constructed from chromatin interaction analysis by pair- end tag sequencing (ChIA-PET), which snapshots chromosomal regions in close proximities mediated by a specific factor [34]. Interestingly, chromosomal contact domains identified by CTCF ChIA-PET data were found to be highly assembled to the TADs identified from Hi-C data [35]. This inspires us to investigate whether putative bovine TADs and CTCF binding motifs could confine bovine-specific regulatory signals, and which is more accurate at defining the regulatory signal. We found both TADs and CTCF binding motifs could predict ASE variation across 18 bovine tissues, and CTCF binding motifs were better at predicting ASE variation than TADs. This finding aligned with previous research from Tang *et al*. [35] who showed that CTCF was responsible for mediating the sub-domain structures within TAD.

To illustrate this, we showed a region on bovine chromosome 3 spanning from position 54 million to position 55.2 million (S1 Figure A). This region was predicted to have frequent internal interactions by all our bovine TAD sets that were mapped from Dixon *et al*. [14]. There were also several CTCF binding motifs found within this region. The genomic regions that were defined only by TAD, only by the more restrictive CTCF binding motifs, and by both TAD and more restrictive CTCF binding motifs all showed significant ASE effects. Inside this 1Mb region, gene *GBP5* (ENSBTAG00000015060), *ENSBTAG00000017670* and *ENSBTAG00000038938* were reported by Chamberlain *et al*. [27] as maternally expressed in white blood cells, and gene *ENSBTAG00000014857, ENSBTAG00000037490* and *ENSBTAG00000038500* appeared to be slightly paternally biased. A cluster of ASE SNPs inside gene *GBP5, ENSBTAG00000017670* and *ENSBTAG00000038938* was significantly maternal-biased. As a result, both our TAD only and CTCF only ANOVA models found this region maternally biased (for this animal at least). Interestingly, gene *GBP5, ENSBTAG00000017670* and *ENSBTAG00000038938* were also located between several CTCF binding motifs. As a result, our TAD+CTCF model found that the CTCF gap maternally biased, and the remaining regions outside CTCF gaps but inside TAD appeared to be paternally biased (S1 Figure). Two other examples of runs of genes which were better predicted by CTCF than by TAD are in provided in S1 Figure B and C.

One explanation why CTCF is better than TAD at predicting ASE regions is that the CTCF binding motifs are within (or close to) the true CTCF binding sequence near the gene that shows significant ASE effect. Since the nucleotide compositions are different between the CTCF binding sequences on parental chromosomes, the CTCF protein preferably binds to one chromosome which resulted in the gene nearby differentially expressed between the parental chromosomes [15, 35]. To test this hypothesis, we examined whether significant aseQTLs and eQTLs were enriched at putative bovine CTCF binding motifs. We found the significant aseQTLs from both bovine white blood and milk cell samples highly enriched at putative bovine CTCF binding motifs, and the significant eQTLs were also highly enriched, though to a less extent than aseQTLs (Table 3; Fig 4). Our results aligned with previous findings from Maurano *et al*. [36], who reported that the variants associated with allelic-imbalanced expression were strikingly concentrated at key positions within CTCF binding motifs, with the higher accessibility allele qualitatively matching the consensus sequence. Our results are also consistent with those from Gómez-Díaz *et al*. [37], who reported that the CTCF protein mediated mono-allelic expression in neuronal differentiation.

Regulatory units are used as a scaffold for testing long-range association or supporting molecular QTLs discovered in humans [35, 38]. To assess whether our predicted bovine TADs have the potential to restrictive hypothesis testing to sites of physical proximity, we examined whether the most significant aseQTL or eQTL for each eGene was more often located in the same TAD as the eGene than not. Our result showed that the furthermost and most-significant white blood and milk cells aseQTL and eQTL were both more likely to be within the same TAD as the eGene than not (Table 4; S4 Table). This indicates that our results could be very useful for reducing the search space for causative regulatory variants and their eGenes. Without knowing the regulatory units, an aseQTL study, which measured ASE at 291,638 genic positions and then tested their association with all SNPs (as identified by the 1000 Bull Genomes Project [1]) within ±1Mb performed 4.8 billion tests [28], and an eQTL study, which tested the association of 11,577 genes and all SNPs within ±1Mb performed 83 million tests [28]. The extremely large number of tests was required because without knowing the regulatory units, for every position or gene, all the SNPs within ±1Mb of the gene had to be tested. If regulatory units were known, the number of association tests could be largely reduced to only test those that were within the same regulatory unit, independent of the distance between the heterozygous SNP and the gene. The number of tests could continue to be reduced by testing associations only between heterozygous SNPs in putative regulatory regions such as enhancers, promoters and CTCF binding motifs. Another benefit of annotating functional regions like TADs and CTCF binding motifs is that new methods that aim to improve the accuracy and reliability of genomic prediction can be devised. Existing methods, such as MultiBLUP [39] and BayesRC [40], could demonstrate higher power by assigning functional classes with potential for a different distribution of effect sizes based on regulatory units. This annotation of regulatory units could also be used with association studies to help prioritise ‘lead’ SNPs.

This study shows that homologous topological association domains (TADs) and CTCF binding motifs can reduce the search space for causative regulatory variants in the bovine genome. Along with study that predict functional elements such as enhancers and promoters across species [41], these results complement the experimental validation that the Functional Annotation of Animal Genomes (FAANG) consortium [42] will produce, accelerating the identification of mutations that affect complex traits.

## Materials and Methods

### Mapping mammalian topological association domains to the bovine genome

Topological association domain (TAD) genomic coordinates from human and mouse embryonic stem cells (hESCs and mESCs) [14], human IMR90 fibroblasts (IMR90) [14] and mouse cortex [32] were obtained from Yue Lab (http://promoter.bx.psu.edu/hi-c/download.html). TAD genomic coordinates from the fresh or frozen liver of mouse and dog were provided as supplementary Table S1 by Rudan *et al*. [16]. Genomic coordinate conversion files, mammalian reference genomes and mammalian genome annotations were all loaded through Bioconductor (version 3.2) [43].

All genomic coordinates of mammalian TADs were converted to the bovine reference genome Bos taurus UMD3.1 (UCSC Genome Browser assembly ID: bostau6) using UCSC Batch Coordinate Conversion program (liftOver) [44] (default settings), which was run inside R (version 3.2.4) using Bioconductor (version 3.2) [43] (AnnotationHub [45], rtracklayer [46] and GenomicRanges [47] packages). Genomic coordinate conversion files were required as an input to liftOver, but were absent for the conversion of some versions of reference genome assemblies. In those cases, an intermediate genome was used to convert the source TADs to bostau6. As a result, the hESC and IMR90 TADs were mapped from hg18 to bostau6 in two steps through an hg19 intermediate. The mESC and mouse cortex TADs were mapped directly to bostau6. The mouse liver TADs were mapped from mm10 to bostau6 through bostau7 (NCBI ID: Btau4.6.1.). The dog liver TADs were mapped to bostau6 from canfam3 through canfam2.

Converting large genomic segments inevitably created a large number of genomic fragments. We employed a recovery procedure, a filtering procedure, and a local refinement procedure to finalise the putative bovine TADs. Throughout this study, we specified “input” as the data that was input to the first step of the genomic conversion, “query” as the data that was input to each step of the genomic conversion, and “target” as the output from each step of the genomic conversion. The recovery procedure was employed in each step of the genomic conversion as follows:

For each input TAD set, *w* was denoted as the minimum width of input TADs.

1. If TAD fragments from the same query sequence were no more than *w* away from each other in the target genome, the TAD fragments were merged into a single TAD in the target genome.
2. If TAD fragments from different query sequences overlapped in the target genome, the TAD fragments were kept as separated fragments in the target genome

The mapping in combination with the recovery procedure resulted in 6 sets of putative bovine TADs (stage 1) from the 6 sets of input TADs respectively. A filtering procedure was then applied to the putative bovine TADs (stage 1) as follows:

1. For each genomic position *i* in the bovine genome, the number of TAD sets that agreed *i* was in a TAD was calculated and denoted as *a_i_*.
2. If a putative bovine TAD (stage 1) was no less than 200 Kb wide, and contained at least one position *i* where *a_i_* was no less than 4, this TAD (stage 1) was kept in the TAD set (stage 2). The 200-Kb threshold was selected because it was able to effectively filter out the small homological genomic fragments which were incapable of forming a TAD in the bovine genome.

The filtering procedure resulted in 6 sets of putative bovine TADs (stage 2) from the 6 sets of putative bovine TADs (stage 1) respectively. A local refinement procedure was then applied to merge any putative bovine TADs (stage 2) that overlapped more than 1 bp. The 1 bp threshold (exclusively) was used, because TADs were calculated from frequencies of interactions between genomic bins [14, 16]. This resulted in the end genomic coordinates for some TADs being the same as the start genomic coordinates for their downstream TAD. These ‘TAD1.TAD2’ structures are maintained in bovine genome, and need to be kept in the final bovine TAD set. However, due to genomic rearrangements, some source TADs from the same TAD set can be mapped to the same location in the bovine genome. Each of these TAD fragments is >200 Kb genomic regions with high frequency of internal interactions. They overlap each other to a great extent in the bovine genome, indicating that they could form into a larger interaction domain, and therefore were merged into one TAD in the bovine genome. The local refinement procedure resulted in 6 sets of putative bovine TADs (final) from the 6 sets of putative bovine TADs (stage 2) respectively.

### Scanning for putative bovine liver CTCF binding motifs

CTCF binding genomic coordinates from the liver tissue of human, mouse, dog and macaque were downloaded from E-MTAB-437 [25]. Mammalian CTCF binding sequences were extracted from masked reference genome hg19, mm9, canfam2 and rhemac2 respectively. Masked reference genome is reference genome where interspersed repeats and low complexity DNA regions are detected and concealed, and is required for motif discovery.

All mammalian CTCF binding sequences were provided as one input to MEME-ChIP [48] in order to identify evolutionarily conserved CTCF binding motifs. Parameter settings for MEME-ChIP were based on the MEME program settings in Vietri Rudan *et al*. [16], who also used the same CTCF ChIP-Seq data from Schmidt *et al*. [25]. We looked for 0 or 1 DNA motif per ChIP-Seq peak, and the maximum input dataset size was adjusted to 70 Mb to account for our input file size.

The latest version of JASPAR CORE motif databases (2016), JASPAR CNE motif database (2008), JASPAR POLII database (2008), JASPAR PHYLOFACTS (2008), UniPROBE [49], and *Homo sapiens* comprehensive model collection database (HOCOMOCO; version 10) were obtained from MEME Suite [50] (http://meme-suite.org/). Transcriptional factor binding motifs that were published to these databases were used to validate our CTCF binding motifs from MEME-ChIP.

Since only the more restrictive set of CTCF binding motifs are reported to transcription factor binding motifs databases, and an increasing number of motifs are discovered as more CTCF ChIP-Seq data are made available, all motifs from MEME-ChIP were input to FIMO [51] (default settings) to scan for putative CTCF binding motifs in the bovine genome. We created two sets of putative bovine CTCF binding motifs from FIMO outputs which were differed by P-value stringencies. Set 1 was a less restrictive set of putative bovine CTCF binding motifs was selected by motif P-value no larger than 10^−5^. Set 2 was a more restrictive set of putative bovine CTCF binding motifs was selected by motif score no less than 80 and motif P-value no larger than 10^−8^ and where overlapping regions were merged. Both sets of putative bovine CTCF binding motifs were used for all the analyses below unless otherwise noted.

### Enrichment of biological hallmarks at topological association domain boundaries

Bovine transfer RNA (tRNA) genes and short interspersed element (SINE) annotations were all loaded through Bioconductor (version 3.2) [43]. Human housekeeping gene names [52] were downloaded as per paper and were used as the list of bovine housekeeping genes. Bovine genomic regions that were enriched for tri-methylation of lysine 4 on histone H3 (H3K4me3) and acetylated lysine 27 on histone H3 (H3K27ac) signals from bovine liver tissue were downloaded from E-MTAB-2633 [53]. Bovine genomic regions that were enriched for H3K4me3 signal from the tender and tough longissimus dorsi of Aberdeen Angus steers were downloaded from GSE61936 [54]. Putative bovine enhancers in homology with human and mouse enhancer databases from VISTA, FANTOM5 and dbSUPER were as described by Wang *et al*. [55].

Putative bovine topological association domain (TAD) boundaries were defined as those genomic intervals, no larger than 400 Kb, between adjacent putative bovine TADs from the same TAD set. For those putative bovine TADs where the end position of the previous TAD was the same as the start position of the following, i.e. a knot, a 400-Kb genomic intervals centring on the “knot” was selected as a TAD boundary. The level of enrichment of the following biological hallmarks was tested at the boundaries from each set of putative bovine TADs: human housekeeping genes, bovine tRNA genes, bovine SINEs, putative bovine CTCF binding motifs, putative bovine enhancer regions in homology with those in VISTA, FANTOM5 and dbSUPER databases, bovine liver H3K4me3 regions, bovine liver H3K27ac regions, bovine tender and tough muscle H3K4me3 regions. The enrichment analysis was run as a permutation test with 10,000 random repeats to test whether the biological signal overlapped significantly more with the putative bovine TAD boundaries than the rest of the genome. We denoted *n* as the number base pairs of a biological signal that overlapped with the putative bovine TAD boundaries in the original dataset.

In the permutation test, for each chromosome, we slid the TAD boundaries along the chromosome by *M* positions, where *M* was a number randomly selected between 1 and the length of the chromosome. After sliding, if a TAD boundary exceeded the range of the chromosome, we recycled the exceeding subset of the TAD boundaries to the start of the chromosome. Then we counted the number of base pairs of a biological signal overlapped with the slid putative bovine TAD boundaries and denoted this number as *m*. The fold change of enrichment was defined as the ratio of *n* to the mean of all *m* values in the 10,000 permutations. The ranking position of *n* within the distribution of all *m* values over all random samples, denoted as *R*, was determined, and a P-value to test the significance of the ranking was computed. For the largest *n* among all *m* values, the P-value was set to < 0.0001 and otherwise it was 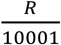. We declared a biological signal “highly enriched” in a set of putative bovine TAD boundaries if *n* was larger than 95% of all *m* values. Our permutation tests resulted in 66 independent analyses (6 sets of finalised putative bovine TADs × 11 biological signals).

### Testing allele-specific expression within regulatory units

Allele-specific expression (ASE) data from 1 lactating cow’s 18 tissues were obtained from Chamberlain *et al*. [27]. The data was the number of mRNA transcripts aligned to each parental allele at all heterozygous exonic positions.

An allele-specific expression (ASE) score was defined as:

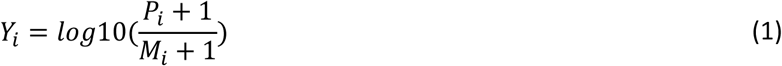
where *i* was a heterozygous SNP in the exon; *Y* was the ASE score from a tissue; *P* was the number of RNA sequence reads from a tissue aligning to position *i* that had the paternal allele; and *M* was the number of RNA sequence reads from a tissue aligning to position *i* that had the maternal allele. One was added to all read counts in each tissue in order to obtain valid ASE scores even for the mono-allelic expressed SNPs, i.e. where only one of the two alleles were expressed.

To examine whether any regulatory units encompassed runs of allele-specifically expressed genes that were observed by Chamberlain *et al*. [27], three independent analyses of variance (ANOVA) models were used. The ASE scores in a tissue were fitted as the response in all three ANOVA models, but the regulatory units in each model were different:
- Model (1) was a TAD only model that defined a regulatory unit only by the putative bovine TADs from a TAD set.
- Model (2) was a CTCF only model that defined a regulatory unit only by the CTCF gaps, which were the genomic intervals between the set 2 CTCF binding motifs (motif score ≥ 80 and motif P-value ≤ 10^−8^).
- Model (3) was a TAD+CTCF model that defined a regulatory unit by both TAD and CTCF gaps. The CTCF gaps were embedded within the putative bovine TADs from a TAD set. Model (3) tested TAD or CTCF gaps accounted for more variation observed in the ASE data.

In each ANOVA model, all innermost genomic regions were required to contain no less than 5 SNPs that had valid ASE scores. Majority of the valid ASE SNPs (>94% in TAD only model, >82% in CTCF only model, and >73% in TAD+CTCF model) were distributed among different genes within the same regulatory unit. If runs of genes within the same regulatory unit did not show any ASE, the mean ASE score of all SNPs within the regulatory unit would be around 0. Alternatively, a paternally-biased regulatory unit would have a mean ASE score larger than 0, and a maternally-biased regulatory unit would have a mean ASE score smaller than 0. There were 72 independent ANOVA tests performed in model (1) which were from the combination of 6 sets of finalised putative bovine TADs and 18 bovine tissues, 18 independent ANOVA tests performed in model (2) from the combination of 1 set of CTCF gaps and 18 tissues, and 72 independent ANOVA tests in model (3) from the combination of 6 sets of finalised putative bovine TADs and 18 tissues. The significance of an ANOVA test was declared at P-value ≤ 10^−8^.

The permutation tests, with 10,000 repeats, were performed to test whether the observed ANOVA result in each model was random. In each permutation test, the ASE scores were shuffled across the whole genome and then the model was refitted with the permuted dataset. The *R*-squared value from the original dataset, *R*, was compared with the 10,000 null R-squared values from the random shuffles, *R*’. A significant ANOVA cohort was declared if *R* was larger than all *R*’ values. In a significant ANOVA test, a regulatory unit was declared to display significant ASE effects if the absolute value of the averaged ASE scores in the region, |*n*|, was larger than 99% of the absolute value of the averaged ASE scores in the permutations |*m*|, i.e. false discovery rate (FDR) < 0.01, and the p-value for *n* was no larger than 10^6−^. There were 72 independent permutation tests performed in model (1), 18 permutation tests performed in model (2) and 72 permutation tests performed in model (3).

### Enrichment of significant aseQTLs and eQTLs within putative bovine CTCF binding motifs

The heterozygous quantitative trait loci (QTLs) that were associated with allelic-specific expression (aseQTLs) and expression variation (eQTLs), were obtained from Chamberlain *et al*. [28]. The aseQTL and eQTL mappings were performed using mRNA transcripts from 141 lactating cows’ white blood and milk cells. The aseQTL and eQTL data was a table of effect and P-value between an aseQTL/eQTL and its eGene target. The RNA sequence data is available from NCBI Sequence Read Archive (Bioproject accession PRJNA305942).

We hypothesized that significant *cis*-expressed QTLs were enriched at putative bovine CTCF binding motifs. We tested this hypothesis using aseQTLs and eQTLs from white blood and milk cells, both of which indicate *cis*- expression QTLs. The significant *cis*-QTLs were defined by a P-value threshold *p*, which was tested at 10^−5^ to 10^−8^. In each significant threshold, we selected a type of *cis*-QTLs (e.g. aseQTLs from white blood cells) whose P-value was no larger than p. In the case where multiple eGene were found to be significantly associated with the same *cis*-QTL, the *cis*-QTL was only counted once. Then we calculated the number of significant *cis*-QTLs within putative bovine CTCF binding motifs, and denoted this number as *n*.

To test the frequency of significant *cis*-QTLs occurring in the rest of the genome, for each chromosome, we slid the significant *cis*-QTLs by *M* positions, where *M* was a number randomly selected between 1 and the length of the chromosome. After sliding, if the position of a *cis*-QTL exceeded the range of the chromosome, we redefined the position of the *cis*-QTL as the slid position subtracting the chromosome length. We denoted *m* as the number of those slid *cis*-QTLs within putative bovine CTCF binding motifs. The slide and recalculation were repeated 10,000 times. The fold change of enrichment was defined as the ratio of *n* to the mean of all *m* values from the 10,000 permutations. The ranking position of *n* within the distribution of all *m* values, denoted as *R*, was determined, and a P-value to test the significance of the ranking was computed. For the largest *n* among all *m* values, the P-value was set to < 0.0001 and otherwise it was 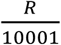. We declared significant *cis*-QTLs highly enriched at putative bovine CTCF binding motifs if *n* was larger than 99% of all *m* values (FDR < 0.01). Our tests resulted in 32 independent analyses (2 types of *cis*-QTLs X 2 cell types × 4 significant thresholds × 2 sets of CTCF binding motifs).

### Testing frequency of the most significant aseQTL or eQTL within the same TAD as eGene

We hypothesized that the most significant *cis*-expressed QTLs for each eGene fell more often within the same TAD as the eGene than not. We tested this hypothesis using aseQTLs and eQTLs, both of which indicate *cis*-expression QTLs. Our *cis*-QTLs were detected using mRNA transcripts from 141 lactating cows’ white blood and milk cells [28]. We selected the most-significant *cis*-QTL for each eGene within a TAD. The most-significant *cis*-QTL for each eGene was required to pass a significance threshold which was tested from 10^−5^ to 10^−8^. In the case where multiple *cis*-QTLs were ranked with the most significance level towards the same eGene, we broke the tie by selecting the *cis*-QTL that was linearly the furthest away from the eGene. The linearly furthermost *cis*-QTL was chosen in order to avoid bias favouring the most significant *cis*-QTL to be within the same TAD as eGene. The result of this procedure was a number of *cis*-QTL and eGene targets, which occurred in the same TAD out of all possible pairs. The expected number of significant *cis*-QTL and eGene within the same TAD was defined as follows:

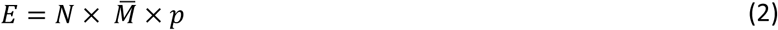
where *E* is the expected number of significant *cis*-QTL and eGene within the same TAD, *N* is the total number of significant eGenes in a TAD that pass the significant threshold *p*, and 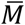 is across chromosome 1 to 29 the maximum number of eSNPs in a chromosome that were used for an aseQTL or eQTL mapping.

A chi-square test was performed to test whether the observed number of furthermost and most-significant *cis*-QTL and eGene within the same TAD was statistically significant or not. The chi-square test was defined as follows:

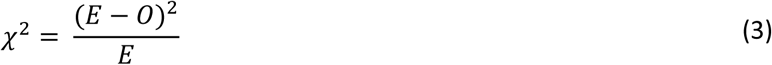
where χ^2^ is the chi-squared value, *E* is defined as above, and *0* is the observed number of significant *cis*-QTL and eGene within the same TAD. A significant chi-squared test was defined as a corresponding chi-square test P-value ≤ 0. 001. There were 96 independent chi-squared tests (2 types of *cis*-QTLs × 2 cell types × 4 significant thresholds × 6 sets of finalised putative bovine TADs).

## List of Abbreviations

ANOVA: Analysis of variance
ASE: allele-specific expression
aseQTL: allele-specific expression quantitative trait loci
ENCODE: Encyclopedia of DNA Elements
eQTL: expression quantitative trait loci
ChIP-Seq: chromatin immunoprecipitation (ChIP) with sequencing
bp: base pair
CTCF: CCCTC-Binding factor
ECDF: empirical cumulative distribution
Hi-C: High-throughput chromosome conformation capture
LD: linkage disequilibrium
TAD: topological association domain
tRNA: transfer RNA
SD: standard deviation
SINE: short interspersed nuclear element
SNP: single nucleotide polymorphism

## Declarations

### Ethics approval and consent to participate

No animal experiments were performed specifically for this work. For the data that were obtained from existing sources, references for these experiments are provided.

### Consent for publication

The authors agree for the publication of this manuscript to BMC Genomics.

### Availability of data and material

Programs, scripts and information for setting up the analysis can be obtained from the authors upon request.

### Author’s contributions

MW performed all the data analysis, conceived the CTCF and ASE study and manuscript preparation. TPH supervised computational and statistical analysis and manuscript preparation. AJC conceived the TAD and ASE study and provided guidance in the ASE study. CJVJ provided the ASE data and guidance in the ASE study. MG designed all enrichment analyses in this study. BGC and JEP assisted with the design of the study and coordination. BJH designed, coordinated and supervised the study and manuscript precreation. All authors read, commented and approved the final manuscript.

## Acknowledgements

We gratefully thank David U. Gorkin (University of California San Diego) for the provision and explanation of TAD data from Dixon *et al*. (2012) study through personal communications.

## Supporting information

### S1 Appendix

**Summary files from MEME-ChIP and FIMO**. Summary file from MEME-ChIP program shows that 4 out of 82 motif clusters are validated in public databases of known transcription factor binding motifs, and the rest are novel transcription factor binding motifs. Summary file from FIMO program shows that the matched CTCF binding motif patterns in the bovine genome.

### S1 Figure

**Runs of genes with allele-specific expression within regulatory units**. The ASE scores (y-axis) for heterozygous loci (x-axis) are plotted as black dot points in each Manhattan plot. The heterozygous locus where ASE score larger than 0 favoured paternal expression and ASE score less than 0 favours maternal expression. The putative bovine TAD is represented as a light pink rectangle centring at y=0. The CTCF binding motifs (motif score ≥ 80 and motif P-value ≤ 10^−8^) are represented as the start and end position of each arrowed curve, where the direction of the arrow is the direction of transcription. Genes are represented as coloured bars starting from y=0 towards either the top (gene on forward strand) or the bottom (gene on reverse strand) of the graph. Gene names are listed in legend and where gene names are not assigned in Ensembl UMD3.1 (bostau6) annotation (release 75), Ensembl gene ID are listed. (A) A region in chromosome 3 is mapped as bovine TAD from human embryonic stem cells, and along with ASE scores from the white blood cells are shown. The ASE SNPs in gene *ENSBTAG00000037490* and *ENSBTAG00000038500* were also within a CTCF gap, but because our ANOVA model required the CTCF gap to contain at least 5 valid ASE SNPs, the CTCF gap that encompassed gene *ENSBTAG00000037490* and *ENSBTAG00000038500* were excluded from TAD+CTCF ANOVA model. (B) A region in chromosome 2 is mapped as bovine TAD from mouse embryonic stem cells, and the ASE scores are from brain caudal lobe. (C) A region from chromosome 23 is mapped as bovine TAD from mouse liver, and the ASE scores are from spleen

### S1 Table

**Summary of genomic conversion of mammalian TADs to the bovine genome.** This table summarises results of converting mammalian TAD coordinates to the bovine genome in each step. Query: the input to liftOver in every step of genomic conversion. Target: the output in every step of genomic conversion. NA: not applicable. Gap: the genomic interval that are not marked as TAD in the data.

### S2 Table

**Rank and fold change of enrichment of biological hallmarks at putative bovine TAD boundaries.** The level of enrichment for each biological signal is measured by the number of base pairs that a biological signal overlapping with the TAD boundary. Two types of TAD boundaries for each TAD set are shown, which are differed by whether a boundary is considered present if two neighbouring putative bovine TADs overlap each other by 1 bp. (A) The degrees of enrichment of all biological signals at the boundary of each TAD set are shown. (B) The degrees of enrichment of each pattern of CTCF binding motifs at the boundary of each TAD set are shown. Each motif pattern has a unique motif number defined by the MEME-ChIP output. Each best possible match motif is manually compared with the MEME-ChIP to access whether the matched motif is previously reported.

### S3 Table

**Enrichment of significant aseQTLs and eQTLs within bovine biological signals**

### S4 Table

**eGenes are often located within the same TAD as their furthermost and most-significant aseQTL/eQTL**

